# Macrophage DYRK1A is broadly required for bacterial phagocytosis

**DOI:** 10.64898/2026.10.09.758040

**Authors:** Madison E. Sanchez, Maureen K. Thomason, Joshua J. Woodward, Michelle L. Reniere

## Abstract

*Listeria monocytogenes* is an intracellular foodborne pathogen that causes the serious illness listeriosis. Macrophages are central to the outcome of *L. monocytogenes* infection, yet the host requirements that facilitate bacterial uptake remain incompletely understood. In a previous genome-wide screen for macrophage factors that influence *L. monocytogenes* infection, we identified the macrophage kinase DYRK1A, which has been studied largely in neurodevelopment and cancer models but has no described role in bacterial infection. Here we show that DYRK1A promotes efficient internalization of *L. monocytogenes* by macrophages. Using several complementary approaches, we find that DYRK1A acts early during infection, primarily by promoting bacterial attachment to the macrophage surface. Transcriptomics of DYRK1A-deficient macrophages revealed reduced abundance of transcripts encoding membrane-associated proteins and components of innate immune and phagocytic pathways, suggesting a transcriptional basis for the attachment defect. Consistent with the broad nature of these changes, the role of DYRK1A extended beyond uptake of *L. monocytogenes* to the Fc receptor pathway and to additional bacterial species spanning multiple genera, establishing a broad role in macrophage phagocytosis. Together, these findings define a previously undescribed role for the host factor DYRK1A in macrophage phagocytosis and expand the function of DYRK1A beyond its established roles to innate immune defense.

## INTRODUCTION

Interactions between bacterial pathogens and professional phagocytes are critical events during all bacterial infections. Phagocyte recognition of an invading pathogen triggers proinflammatory signaling and cytokine release, priming the immune system for a robust infection response (1). Simultaneously, the cell undergoes cytoskeletal rearrangement to engulf the invading bacterium (2,3). Phagocytes such as macrophages subject internalized bacteria to an array of antimicrobial mechanisms, including digestive enzymes, reactive oxygen and nitrogen species, and acidification, while bacterial pathogens use specific virulence factors to subvert these immune attacks and, in some cases, establish replicative niches that promote infection (4,5). While much is understood about the mechanisms of phagocytosis and cytokine release from host phagocytes, less is known about regulatory pathways that may alter or mediate these processes, particularly across the range of pathogen-specific responses.

*Listeria monocytogenes* is the etiologic agent of listeriosis, a rare but serious foodborne illness causing mortality rates of ∼20% among hospitalized patients (6). *L. monocytogenes* is one of only three FDA-designated zero-tolerance pathogens and was the primary cause of microbial-related food recalls in the US and Canada between 2012 and 2022 (7,8). This dual burden makes it a major economic and public health concern. *L. monocytogenes* is also a model pathogen in the research setting, where its intracellular lifecycle has been defined and studied for over 35 years (9). This lifecycle requires the recognition and internalization of *L. monocytogenes,* followed by its escape from the vacuole into the cytoplasm where bacterial replication and cell-to-cell spread occur (9). This general lifecycle is well-established, and more recently, a growing body of literature has begun to reveal host cell-type- and infection-site-specific nuances that refine the original model (10–13). For example, *L. monocytogenes* is internalized by both phagocytic and non-phagocytic cells, though the receptor-ligand interactions and signaling underlying its invasion of non-phagocytic cells are better defined than those enabling its uptake by phagocytic cells (14). Phagocytic cells, including macrophages, play a dual role during *L. monocytogenes* infection by either promoting infection and dissemination when in a growth-permissive state (15,16), or by becoming bactericidal and host-protective later during infection upon activation (17,18). Our current understanding of the host factors mediating phagocytosis focuses primarily on the bactericidal complement-mediated internalization pathway (18–20), while the mechanisms underlying *L. monocytogenes* uptake by growth-permissive macrophages, which propagate infection, remain relatively unexplored.

To identify macrophage factors important during *L. monocytogenes* infection, we previously performed a genome-wide CRISPR/Cas9 screen (10). That study identified 235 host genes important during *L. monocytogenes* infection of macrophages and revealed a regulatory role for the tumor suppressor PTEN (Phosphatase and Tensin Homolog) during phagocytosis. PTEN promotes phagocytosis of *L. monocytogenes* by enhancing adherence of the bacteria to the cell surface. This interaction supports disease progression, as mice lacking PTEN in myeloid cells were unable to restrict *L. monocytogenes* growth during foodborne listeriosis. This discovery demonstrates the power of the screen to identify host factors with meaningful roles during infection. We therefore sought to leverage those data to uncover additional genetic requirements in *L. monocytogenes* infection. Specifically, we became interested in *Dyrk1a*, one of the top ten most significant genes identified and validated in the screen. *Dyrk1a* was also one of only nine genes for which all four sgRNAs were significantly depleted; it also functions independently of the PTEN pathway and has no known role during bacterial infections.

*Dyrk1a* encodes <u>D</u>ual-specificity T<u>y</u>rosine Phosphorylation-regulated <u>K</u>inase 1A (DYRK1A), a kinase that localizes primarily to the nucleus, with a smaller pool active in the cytosol (21–23). It phosphorylates a broad range of substrates, including proteins that regulate transcription, giving it wide-reaching effects on gene expression and cellular regulation (24–26). DYRK1A is dosage-sensitive, as triplication of chromosome 21, which contains DYRK1A, contributes to the neurological features of Down syndrome, while DYRK1A haploinsufficiency causes a distinct syndrome featuring intellectual disability and autism spectrum features (27,28). Due to its pronounced neurological effects, DYRK1A has been studied predominantly in neurons and other brain cell types. Only recently has research expanded to immune cells, primarily B cells and T cells (29,30), and its role in phagocytic cells such as macrophages remains largely unexplored.

In this study, we sought to analyze the role of DYRK1A during distinct stages of the *L. monocytogenes* lifecycle in macrophages. We demonstrate that DYRK1A is required early during macrophage infection for efficient bacterial internalization, specifically for the adherence of *L. monocytogenes* to the macrophage membrane. Using bulk RNA-seq, we identified differences in basal transcript abundance between *Dyrk1a*^−/−^ and control macrophages, revealing changes in transcript levels for genes involved in innate immune signaling and in pathways regulating membrane organization and phagocytic machinery. This transcriptional signature is further supported by our finding that the requirement for DYRK1A during phagocytosis extends beyond *L. monocytogenes* to diverse bacterial species. Overall, this work identifies DYRK1A as a newly recognized, broadly required regulator of macrophage phagocytosis.

## RESULTS

### DYRK1A is required for *Listeria monocytogenes* uptake into macrophages

In our previous genome-wide screen, *Dyrk1a* emerged as one of 235 genes required for efficient *L. monocytogenes* infection of immortalized bone marrow-derived macrophages (iBMMs), with *Dyrk1a*-deficient cells showing the strongest phenotype among those analyzed (10). The screen was performed 6 hours post-infection (p.i.), during which time *L. monocytogenes* attaches to and is internalized by host cells, escapes the vacuole, and begins replicating in the host cytosol within 1 hour. Thus, it remains unclear when and where DYRK1A limits *L. monocytogenes* infection of macrophages.

CRISPR/Cas9 gene editing produced two independent *Dyrk1a*-deficient iBMM cell lines, each generated using a unique sgRNA cut site (10). As each cell line is a heterogeneous population of edited and unedited cells, we used Interference of CRISPR Edits (ICE) analysis, a genetic method for estimating knockout efficiency, to calculate the likelihood of functional *Dyrk1a* expression in each population (31). The *Dyrk1a*-deficient sgRNA 1 and sgRNA 2 iBMM populations have knockout efficiency scores of 84% and 89%, respectively, indicating that most macrophages in the population do not express functional *Dyrk1a*. Immunoblotting corroborated this result, showing no detectable DYRK1A protein in either population (Fig 1A). In contrast, control cells transfected with an empty vector lacking a sgRNA sequence (empty vector, EV) showed a strong DYRK1A band.

**Figure 1.**
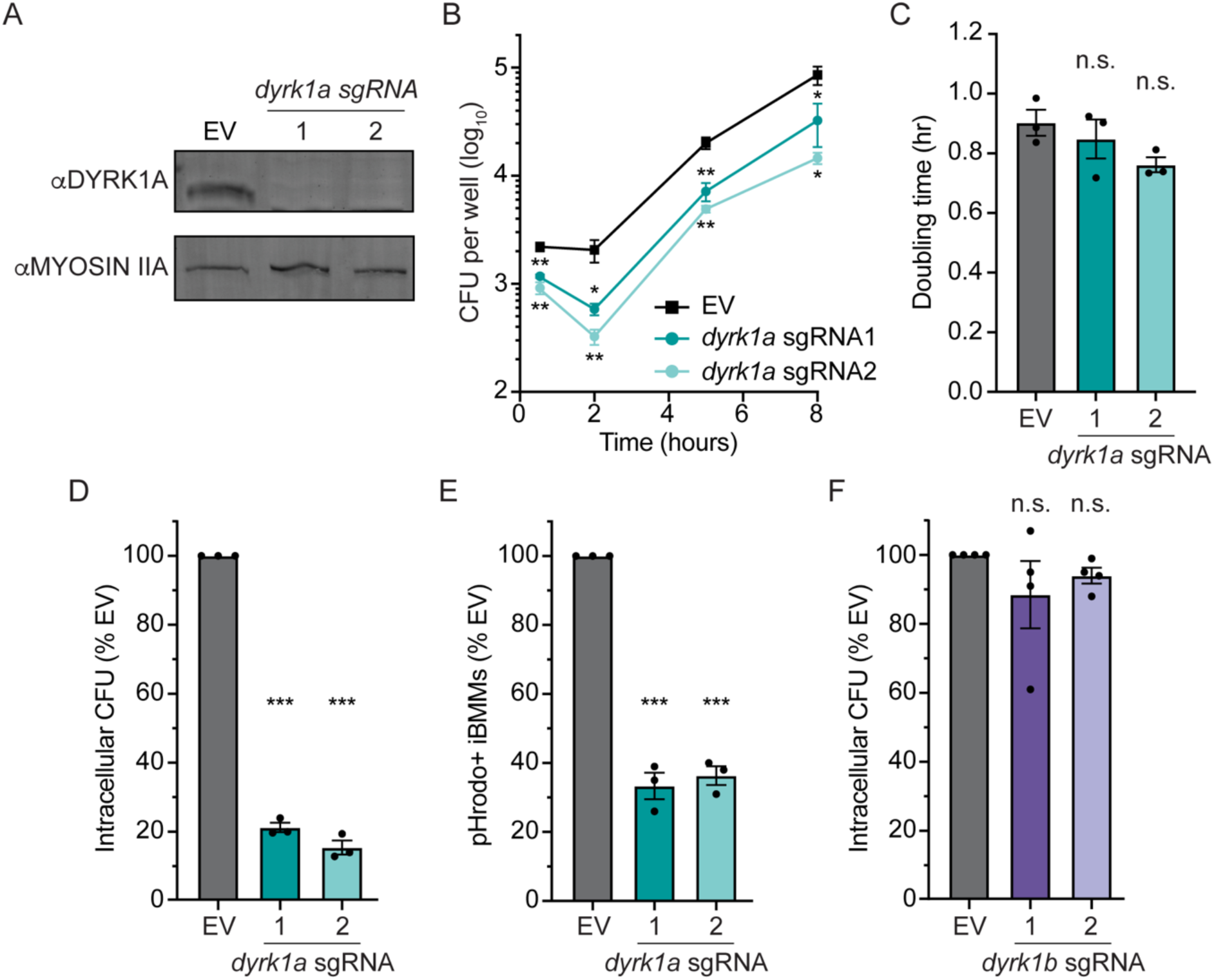
DYRK1A is required during early stages of *L. monocytogenes* infection of iBMMs. **(A)** DYRK1A immunoblot, with Myosin IIA as a loading control. **(B)** Intracellular growth of *L. monocytogenes* in *Dyrk1a*-deficient cell lines. Gentamicin was added 30 min post-infection (p.i.) and CFU were enumerated at each timepoint. **(C)** Quantification of intracellular *L. monocytogenes* doubling time in *B,* 2-5 hours p.i.. **(D)** Gentamicin protection assay quantifying intracellular *L. monocytogenes* CFU 1-hour p.i. in *Dyrk1a*-deficient and EV iBMMs (MOI = 1). **(E)** pHrodo phagocytosis assay quantifying the proportion of iBMMs containing pHrodo-labeled *L. monocytogenes* by flow cytometry at 30 minutes p.i. (MOI = 10). **(F)** Gentamicin protection assay quantifying intracellular *L. monocytogenes* CFU 1-hour p.i. in *Dyrk1b*-deficient and EV iBMMs (MOI = 1). **(C-F)** Each data point is the mean of two technical replicates and bars are means and SEM of at least three biological replicates. **(D-F)** Data are normalized to EV iBMMs. \**p* < 0.05, \*\**p* < 0.01, \*\*\**p* < 0.001, n.s. = non-significant (*p* > 0.05), as determined by Dunnett’s multiple comparisons test compared to EV in each condition.

To begin to identify the stage or stages of the *L. monocytogenes* infection cycle that require DYRK1A, we infected *Dyrk1a*-deficient iBMMs and monitored bacterial internalization and cytosolic replication over time using an intracellular growth curve. In this assay, gentamicin is added to the medium 30 minutes p.i. to kill extracellular bacteria, allowing intracellular *L. monocytogenes* to be quantified at defined time points thereafter. As early as 30 minutes p.i., *Dyrk1a*-deficient cells contained significantly fewer cell-associated and internalized *L. monocytogenes* compared to EV control iBMMs (Fig 1B), indicating that DYRK1A is important during the early stages of the infection cycle. However, there was no difference in the doubling time of intracellular *L. monocytogenes* between 2 and 5 hours p.i. (Fig 1C), suggesting that DYRK1A does not regulate *L. monocytogenes* cytosolic growth. Together, these data indicate that DYRK1A-dependent regulation of *L. monocytogenes* infection occurs during the initial stages of internalization and adherence to the macrophage, rather than during the later stage of cytosolic replication.

To more precisely investigate the role of DYRK1A during the initial stages of infection, we quantified the total number of internalized bacteria across the macrophage population 1 hour p.i. using a gentamicin protection assay. *Dyrk1a*-deficient cell lines were infected with *L. monocytogenes* and briefly centrifuged to synchronize bacterial contact with the macrophages. At 30 minutes p.i., cells were treated with gentamicin-containing medium to kill extracellular bacteria, and intracellular bacteria were quantified by colony forming units (CFU) at 1 hour p.i.. *Dyrk1a*-deficient cell lines exhibited a significant ∼80% decrease in total intracellular CFU compared to control iBMMs (Fig 1D), indicating that DYRK1A is required for internalization of *L. monocytogenes*.

Gentamicin protection assays reflect total bacterial numbers but cannot distinguish whether a few macrophages engulfed many bacteria or many macrophages engulfed one bacterium. We therefore used flow cytometry as a complementary approach to measure the proportion of macrophages that internalized bacteria during early infection. *L. monocytogenes* were conjugated to pHrodo, a pH-sensitive dye that fluoresces upon exposure to an acidified compartment, such as the macrophage phagosome, enabling quantification of bacteria-containing cells by flow cytometry. After 30 minutes of infection, *Dyrk1a*-deficient macrophage populations showed a significant ∼65% decrease in the proportion of bacteria-containing cells (Fig 1E). Together, these results suggest that DYRK1A is required for efficient uptake of *L. monocytogenes* into macrophages during the first hour of infection.

DYRK1A is one of five members of the DYRK protein kinase family and is evolutionarily most similar to DYRK1B, sharing a highly conserved catalytic domain (32–34). Given the similarity, we tested whether the requirement for DYRK1A during *L. monocytogenes* internalization is specific to DYRK1A or shared with DYRK1B. We generated two independent *Dyrk1b*-deficient iBMM cell lines using CRISPR/Cas9 editing, with knockout efficiency scores of 76% and 79% by ICE analysis. Gentamicin protection assays revealed no difference in intracellular *L. monocytogenes* between *Dyrk1b*-deficient cell lines and control cells (Fig 1F). This indicates that, despite their catalytic similarity, *L. monocytogenes* internalization depends specifically on DYRK1A rather than being a shared feature of the DYRK1 kinases.

### Macrophage DYRK1A promotes *L. monocytogenes* adherence and internalization

Early bacterial infection of macrophages can be delineated into two distinct stages: the specific adherence of a bacterium to the host cell and the mechanical re-localization of that bacterium into the phagosome. Apart from our group’s previous studies on the PTEN pathway, little is known about the macrophage requirements for either of these critical processes during *L. monocytogenes* infection. By determining the role of DYRK1A in these pathways, we can begin to dissect the essential host processes required for the early stages of infection in macrophages.

To study DYRK1A in the discrete processes of adherence and internalization, we used immunofluorescence microscopy to differentiate intracellular and extracellular *L. monocytogenes* during infection of macrophages. iBMMs were infected with *L. monocytogenes* constitutively expressing *mCherry* and labeled under non-permeabilizing conditions with α-*Listeria* antisera, then counterstained with AlexaFlour488. Thus, all *L. monocytogenes* were mCherry^+^, while only extracellular bacteria were additionally labeled with AlexaFluor488 (Fig 2A). To ensure analytical rigor, over 2,000 macrophages in each cell line were quantified per replicate. First, adherence was calculated as total bacteria per 100 macrophages. This includes both intracellular and extracellular bacteria, since a bacterium must adhere before internalization. To capture the kinetics of adherence during early infection, we quantified bacteria at 15, 30 and 60 minutes p.i.. *Dyrk1a*-deficient iBMMs had ∼50% fewer adherent bacteria per 100 macrophages when compared to control cells as early as 15 minutes p.i., and this adherence defect persisted throughout the first hour of infection (Fig 2B). These data show that DYRK1A promotes the physical attachment of *L. monocytogenes* to the macrophage membrane, a critical step in the *L. monocytogenes* infection cycle.

**Figure 2.**
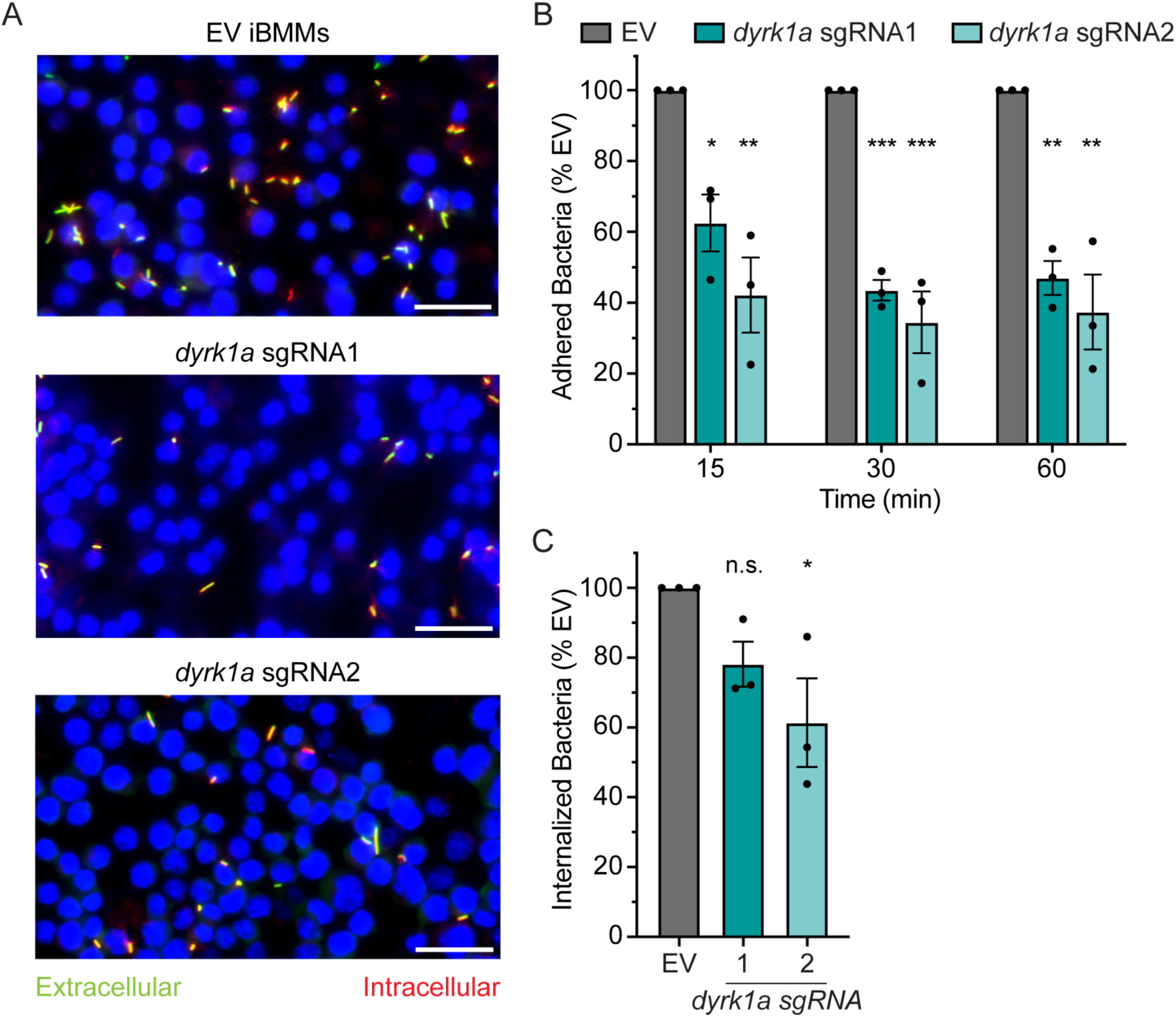
DYRK1A primarily modulates adherence of *L. monocytogenes* to macrophages. **(A)** Representative immunofluorescence microscopy images of *L. monocytogenes* infection assay quantified in *B*, 15 min p.i. (MOI = 5). Extracellular *L. monocytogenes* appear green while intracellular bacteria are red. Nuclei were stained with DAPI (blue). Scale bars represent 25 µm. **(B)** Quantification of *L. monocytogenes* adhered to cells at 15 min p.i. (MOI = 5), 30 min p.i. (MOI = 2) and 60 min p.i. (MOI = 2). Total bacteria per 100 iBMMs were counted for three independent images per cell line (totaling ≥ 2,000 macrophages) and averaged for each replicate. **(C)** Quantification of *L. monocytogenes* internalization at 60 min p.i. (MOI = 2). Total and extracellular bacteria were counted for three independent images per cell line and averaged for each replicate. Internalization was calculated as the percentage of intracellular bacteria (total minus extracellular) relative to total *L. monocytogenes*. **(B-C)** Data are normalized to EV. Mean values of three independent images are represented as individual data points. Data are means and SEM of three biological replicates. \**p* < 0.05, \*\**p* < 0.01, n.s. = non-significant (*p* > 0.05), as determined by Dunnett’s multiple comparisons test compared to EV in each condition.

We next tested whether DYRK1A also affects the mechanical process of phagocytosis after *L. monocytogenes* has attached to the membrane. Using the same fluorescent microscopy approach described above, internalization was calculated as the percentage of total mCherry^+^ bacteria that were AlexaFluor488-negative per image. This calculation only considers bacteria that have already successfully adhered to a macrophage, separating internalization from our previously calculated adherence metrics. Because internalization requires time following adherence, few bacteria are internalized at early timepoints in either genotype. In *Dyrk1a*-deficient cells, this is compounded by fewer adhered bacteria overall, resulting in correspondingly fewer internalization events to quantify. We therefore restricted our internalization analysis to 60 minutes p.i., when sufficient events had accumulated for reliable quantification. At this timepoint, *Dyrk1a*-deficient iBMMs trended toward decreased internalization of *L. monocytogenes* compared to empty vector cells, though this difference only reached statistical significance for one *Dyrk1a*-deficient population (Fig 2C). These results suggest that DYRK1A primarily promotes phagocytosis by enhancing *L. monocytogenes* attachment to macrophages, with more modest effects on subsequent bacterial engulfment.

### DYRK1A shapes the macrophage transcriptional landscape associated with bacterial attachment and phagocytosis

The first physical interaction during infection is the attachment of a bacterium to the host cell surface, which requires a bacterial ligand and its corresponding host membrane receptor. DYRK1A is required for efficient *L. monocytogenes* attachment to macrophages, an interaction likely mediated by an unidentified host receptor. DYRK1A itself is unlikely to serve as this receptor directly, since it localizes to the nucleus and cytosol rather than the cell membrane. However, many DYRK1A substrates are transcription factors that drive broad changes in gene expression. Because the DYRK1A regulon has not been defined in macrophages, we tested whether basal transcriptional differences in *Dyrk1a*-deficient macrophages could point toward candidate pathways underlying the adherence defect.

To identify potential DYRK1A-dependent membrane receptors, we sought to characterize the resting transcriptomes of *Dyrk1a-*deficient macrophages compared to control cells. As bacterial attachment requires the receptor to be present before infection occurs, we focused on basal transcriptional differences between uninfected cell lines. A potential complication is that the mixed *Dyrk1a*-deficient populations retain a small fraction of *Dyrk1a*-expressing cells that could mask smaller transcriptional differences. We therefore performed single-cell fluorescence-activated cell sorting (FACS) of each *Dyrk1a*-deficient population and selected two clonal lines from each, yielding four clonal *Dyrk1a^−/−^* lines in total. Genetic validation by Sanger sequencing confirmed that each clonal line harbored frameshifts or deletions predicted to result in truncation of DYRK1A. Each *Dyrk1a^−/−^* cell line was also phenotypically validated using a gentamicin protection assay (Fig 3A), showing a decrease in intracellular *L. monocytogenes* compared to control cells, similar to the decrease observed in the original *Dyrk1a*-deficient populations.

**Figure 3.**
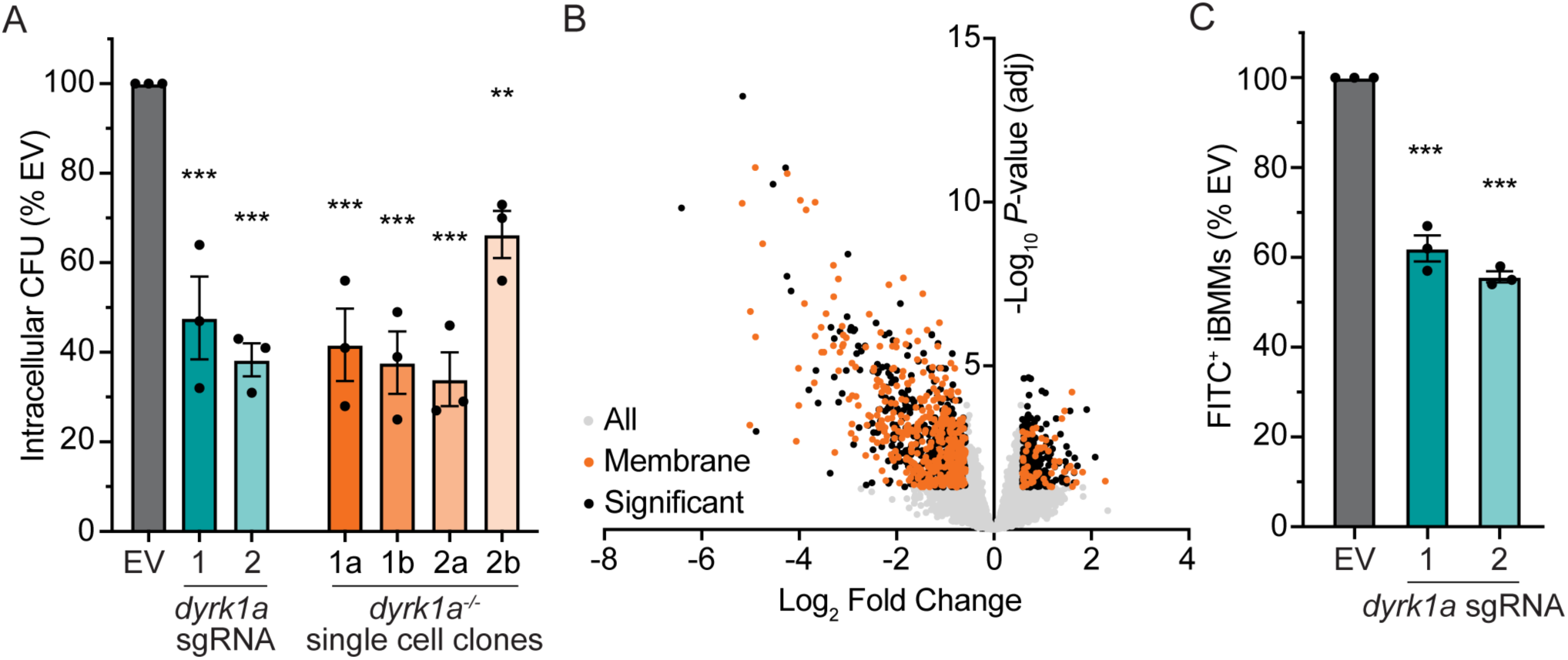
DYRK1A loss alters transcript abundance of genes linked to membrane proteins and innate immunity. **(A)** Gentamicin protection assay comparing *Dyrk1a*-deficient cell lines and *Dyrk1a^−/−^* single cell clones, quantifying intracellular *L. monocytogenes* 1-hour p.i. (MOI = 1). Each data point represents the average of two technical replicates. (**B**) Volcano plot of differentially expressed transcripts identified by RNAseq. Transcripts meeting a significance threshold of FDR < 0.05 and |log_2_ fold-change| > 1.5 are displayed in black; among these, genes encoding membrane proteins are represented in orange. **(C)** Phagocytosis of IgG-opsonized, FITC-labeled beads, measured by flow cytometry 30 minutes after bead addition. **(A,C)** Data are means and SEM of three biological replicates, normalized to EV. \*\**p* < 0.01, \*\*\**p* < 0.001, as determined by Dunnett’s multiple comparisons test compared to EV.

To assess basal transcription levels, RNA was extracted from all four *Dyrk1a^−/−^* and empty vector cell lines 24 hours after seeding, and bulk transcript levels were quantified. Transcripts corresponding to 835 genes were significantly decreased in abundance in all four *Dyrk1a^−/−^* single cell clonal lines compared to empty vector control (S1 Table), and 414 of these genes encode membrane-associated proteins, based on UniProt subcellular location annotations (Fig 3B). While these data do not identify the specific receptor or receptors mediating *L. monocytogenes* attachment, they show that DYRK1A loss broadly influences transcription of the macrophage membrane protein landscape, consistent with the adherence defect we observed.

To more broadly characterize the transcriptional consequences of DYRK1A loss, we performed gene set enrichment analysis (GSEA) and visualized enriched pathways as a network using Cytoscape with the EnrichmentMap plugin (Fig S1A) (35). In this network, nodes represent gene sets and edges connect gene sets that share overlapping genes. Many resulting clusters, including cell cycle, transcription, and translation, correspond to pathways already known to involve DYRK1A in other cell types(36–38), suggesting that many regulatory roles of DYRK1A are conserved in macrophages. Despite this conservation, we observed that some of the pathways we identified are particularly relevant for macrophage function during infection. From these critical pathways, we further identified and categorized genes involved in cytoskeletal rearrangement, inflammatory cytokine signaling (IL-6, TNF, IL-8, Type II IFN, IL-1β), phagocytosis and endocytosis, and innate immune surface receptors (including TLR4) (S2 Table, Fig S1B). Within the phagocytosis and endocytosis grouping, we found that many functional categories contain genes that may be responsible for bacterial attachment and internalization dynamics in macrophages (Table 1). For example, we identified common phagocytic receptors not previously linked to DYRK1A (*CD36* and *Scarb1),* along with genes for extracellular sensing that regulate phagocytosis and surface attachment. Src-kinase and the MAPK signaling alter the cytoskeletal and immune response after receptor clustering. Relatedly, we identified genes involved in actin and cytoskeletal dynamics that influence phagosome formation. These emerging patterns suggest a multifaceted role for DYRK1A, indicating that its loss may affect multiple stages of phagocytosis.

**Table 1.**
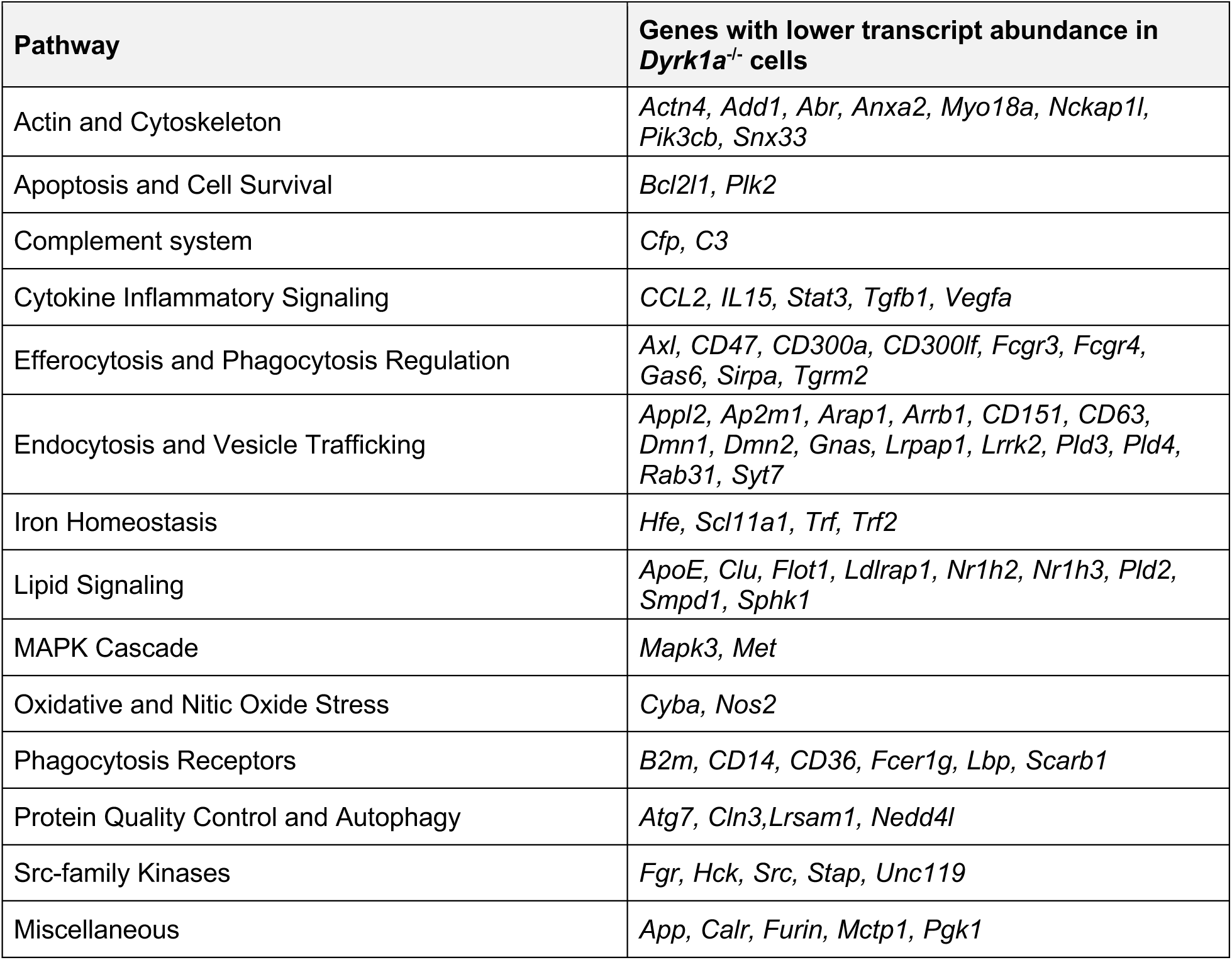
Genes with decreased transcript abundance in *Dyrk1a^−/−^* macrophages from pathways involved in phagocytosis and endocytosis.

During bacterial infection, cell surface receptor recognition, cytoplasmic signaling, and cytoskeletal rearrangement must act together to enable internalization. Because our transcriptomic data showed decreased transcript abundance in many genes involved in each of these processes, we hypothesized that the role of DYRK1A in phagocytosis might extend beyond *L. monocytogenes*-specific internalization. To test this using a receptor pathway distinct from *L. monocytogenes*, we examined phagocytosis mediated by Fc receptors, which recognize IgG-opsonized particles. *Dyrk1a*-deficient macrophages were treated with IgG-opsonized, fluorescein-conjugated latex beads for 30 minutes, and bead uptake was quantified as a measure of Fc receptor-mediated phagocytosis. *Dyrk1a*-deficient macrophages showed a decreased ability to internalize these beads compared to EV control cells (Fig 3C), indicating that DYRK1A is required for efficient Fc receptor-mediated phagocytosis, a pathway unrelated to *L. monocytogenes* recognition. The breadth of phagocytosis, endocytosis, and innate immune genes downregulated in DYRK1A-deficient macrophages parallels this functional result. Together, they indicate that DYRK1A acts as a general regulator of the macrophage phagocytic program rather than a mediator of any single recognition pathway.

### Macrophage DYRK1A is broadly required for bacterial internalization

Given the breadth of membrane and innate immune-related transcripts affected in *Dyrk1a^−/−^* macrophages, together with our finding that DYRK1A is required for phagocytosis independent of *L. monocytogenes* recognition, we hypothesized that DYRK1A may broadly support bacterial internalization. To test whether this role extends beyond a mechanism specific to *L. monocytogenes*, we measured internalization of several additional Gram-positive and a Gram-negative bacterial species in *Dyrk1a*-deficient and EV control macrophages. For each bacterial species, we performed gentamicin protection assays to quantify the total number of internalized bacteria, as well as pHrodo phagocytosis assays to determine the proportion of the macrophage population capable of internalizing one or more bacteria.

Within the *Listeria* genus, we investigated the role of DYRK1A in macrophage internalization of two additional species: *L. innocua and L. ivanovii*. *L. innocua* is a non-pathogenic species closely related to *L. monocytogenes* but lacking the pathogenicity island that harbors canonical *L. monocytogenes* virulence factors, whereas *L. ivanovii* is pathogenic but evolutionarily more distant from *L. monocytogenes*. Interestingly, *Dyrk1a-*deficient iBMMs internalized fewer bacteria than EV control cells for each species tested (Fig 4A). The proportion of macrophages capable of internalizing each species was also reduced relative to EV control macrophages (Fig 4B). These data reveal that DYRK1A is broadly required for early internalization of distinct *Listeria* species by macrophages, and that the role of DYRK1A in internalization is not specific to pathogenic bacteria.

**Figure 4.**
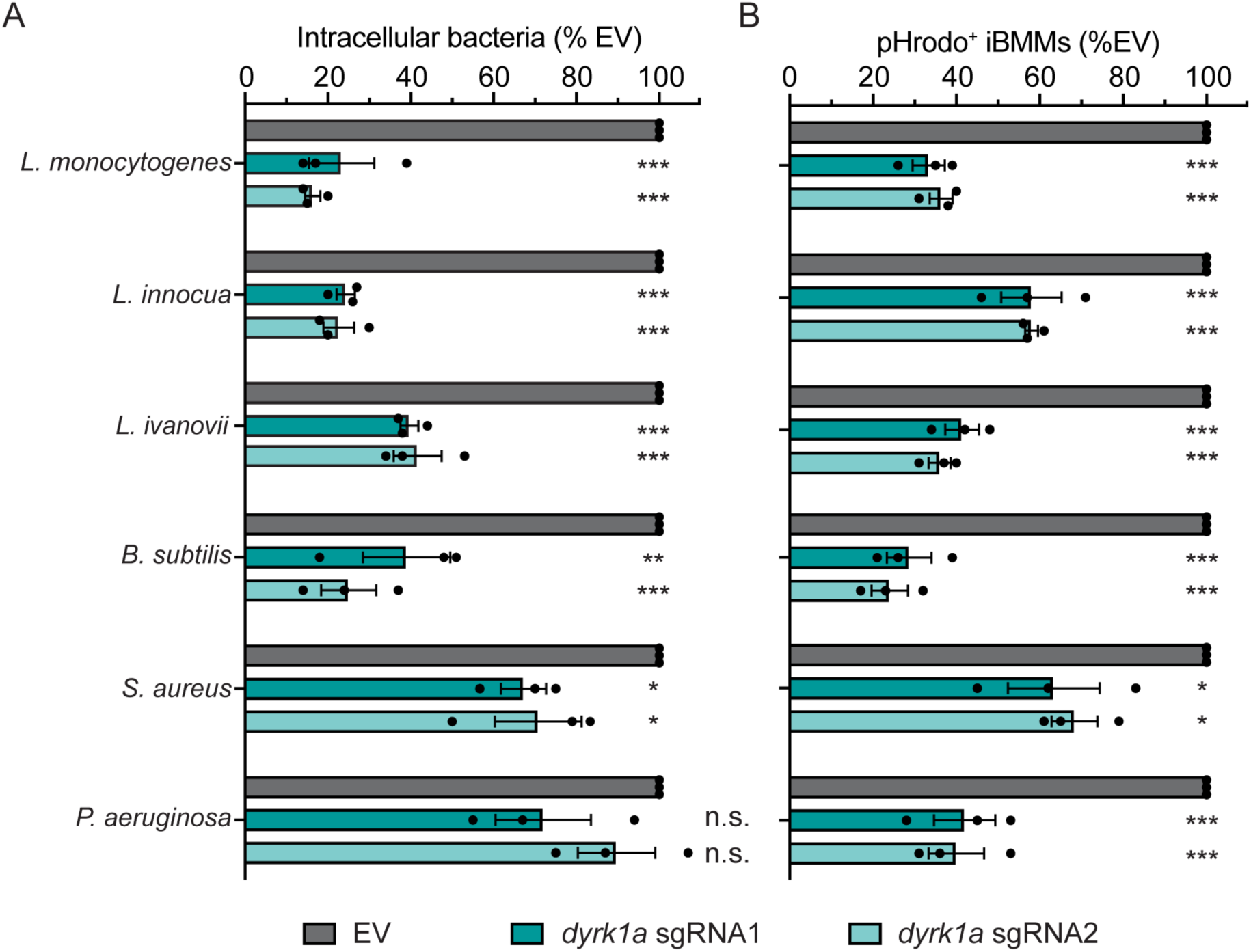
Internalization of diverse bacterial species is impaired in *Dyrk1a-*deficient macrophages. **(A)** Gentamicin protection assay measuring intracellular bacteria 1-hour p.i. in *Dyrk1a*-deficient and EV iBMM cell lines. Species-specific MOIs used: *L. monocytogenes,* 1; *L. innocua,* 2; *L. ivanovii*, 5; *B. subtilis,* 100; *S. aureus* strain JE2, 0.001; *P. aeruginosa,* 10. Each data point represents the mean of two technical replicates. **(B)** pHrodo phagocytosis assay measuring bacterial internalization at 30 minutes p.i.. Cell lines were infected with pHrodo-labeled bacteria at the following MOIs: *Listeria spp.,* 10; *B. subtilis*, 0.1; *S. aureus,* 0.1; *P aeruginosa,* 10. Macrophages that internalized bacteria were quantified by flow cytometry **(A-B)** Data are means and SEM of at least three biological replicates and are normalized to EV iBMM internalization for each species. \**p* < 0.05, \*\**p* < 0.01, \*\*\**p* < 0.001, n.s. = non-significant (p > 0.05), as determined by Dunnett’s multiple comparisons test compared to EV.

We next sought to expand our investigation beyond *Listeria spp*. to explore how broadly DYRK1A is required for bacterial internalization. As a representative panel, we investigated two non-*Listeria* Firmicutes, the non-pathogenic environmental saprophyte *Bacillus subtilis* and the pathogenic *Staphylococcus aureus*, as well as the Gram-negative Gammaproteobacteria *Pseudomonas aeruginosa*. For *B. subtilis*, *Dyrk1a*-deficient macrophages internalized fewer bacteria by gentamicin protection assay, and fewer macrophages internalized bacteria by pHrodo assay, compared to controls (Fig 4A-B). Loss of DYRK1A significantly reduced internalization of *S. aureus* (Fig 4A-B), confirmed for both the laboratory strain Newman and the clinical methicillin-resistant isolate JE2 (Fig S2). For *P. aeruginosa*, the number of bacteria internalized did not differ significantly by gentamicin protection assay (Fig 4A), but significantly fewer macrophages internalized bacteria by pHrodo assay (Fig 4B). These data suggest that DYRK1A broadly supports internalization of bacteria by phagocytes, identifying a previously undescribed role for DYRK1A in myeloid innate immunity.

## DISCUSSION

DYRK1A is a eukaryotic kinase with no previously described role in bacterial infection of macrophages. Here we show that DYRK1A is required for macrophages to internalize *L. monocytogenes*, using complementary approaches that quantify intracellular bacteria and the proportion of macrophages containing internalized bacteria. Microscopic studies localize this requirement to an early step of the *L. monocytogenes* intracellular lifecycle, the attachment to the macrophage surface, with a possible smaller contribution to translocation into the cell. Profiling the transcriptome of uninfected *Dyrk1a^−/−^* macrophages revealed a lower abundance of transcripts encoding membrane-associated proteins and genes for innate immune signaling, cell-surface receptors, actin dynamics, and phagocytosis. The requirement extended beyond *L. monocytogenes* to additional bacteria. Together, these findings establish DYRK1A as a previously unrecognized host factor for *L. monocytogenes* infection and a broad promoter of bacterial phagocytosis. They also identify a role in shaping the membrane and innate immune transcriptome of macrophages, expanding its known functions beyond the nervous system.

Despite the influence of DYRK1A on major biological processes, few studies have considered its role during microbial infection. A small body of work identified DYRK1A as a pro-viral factor for coronaviruses, through transcriptional regulation of the ACE2 or ANPEP receptors, and for cytomegalovirus, through direct phosphorylation of a viral protein (39–41). During HIV-1 infection, DYRK1A instead acts as an antiviral factor, restricting viral transcription and replication in an NFAT-dependent manner (42). In macrophages and microglia stimulated with LPS, DYRK1A promotes TLR4 signaling and pro-inflammatory cytokine release (43,44). Across these examples, DYRK1A can promote or restrict viral infection and shape innate immune signaling, yet its role in bacterial infection had not yet been examined.

Our previous CRISPR screen for macrophage genes that promote *L. monocytogenes* infection identified over 230 genes, with DYRK1A among the most promising candidates (10). That screen captured the *L. monocytogenes* infection cycle only through the stages before cell-to-cell spread and was designed to identify host factors mediating internalization, vacuolar escape, or cytosolic replication. Among these, we find the requirement for DYRK1A is limited to internalization, indicating that distinct macrophage genes act at separable steps of the lifecycle. The same screen identified PTEN, another factor required for *L. monocytogenes* internalization. PTEN promotes uptake of *L. ivanovii* and specific *L. monocytogenes* serovars and acts only at attachment, whereas DYRK1A is required for both *Listeria* and non-*Listeria* species and may also contribute to the later internalization step. This distinction identifies DYRK1A as a broader regulator of phagocytosis than PTEN, and defining how the two pathways intersect will require further work.

We further found that DYRK1A, but not DYRK1B, is required for *L. monocytogenes* internalization. DYRK1A and DYRK1B comprise the class I DYRK subfamily and share a conserved catalytic domain, to the extent that small-molecule DYRK1B inhibitors frequently cross-react with DYRK1A (45). Both kinases also require the same PHD1-dependent prolyl hydroxylation step for activation (46). Despite these similarities, their interactomes diverge, in part because of three residue differences between their catalytic domains (33,34). The specific requirement for DYRK1A therefore indicates that *L. monocytogenes* internalization is not a shared feature of the DYRK family and likely depends on these functional differences. Comparative interactome profiling of DYRK1A and DYRK1B could filter out shared interactions and narrow down which DYRK1A-specific functions drive *L. monocytogenes* phagocytosis.

Given its established roles in the actin cytoskeleton and endocytic machinery, we expected DYRK1A to strongly promote *L. monocytogenes* internalization, so the comparatively modest contribution to this step was unexpected. DYRK1A physically interacts with actin, although it reportedly promotes or prevents polymerization depending on its localization and interacting partner (47–49). It is also required for clathrin-mediated neuronal endocytosis through interactions with clathrin, dynamin 1, and other associated proteins (50,51). In *Dyrk1a^−/−^*macrophages, transcripts for actin and clathrin-mediated endocytic machinery, including those encoding dynamin (*Dnm1* and *Dnm2*), are reduced. This reduction would be expected to impair clathrin-dependent uptake, yet the internalization step was only modestly and inconsistently affected. The expectation that clathrin and dynamin matter comes from non-phagocytic cells, where *L. monocytogenes* invasion depends heavily on both (56). One explanation is that macrophage phagocytosis of *L. monocytogenes* relies less on clathrin-mediated machinery than the invasion of non-phagocytic cells by *L. monocytogenes*. Alternatively, the modest internalization phenotype may reflect the limited sensitivity of that measurement rather than true clathrin independence. Nevertheless, our results suggest that the dominant contribution of DYRK1A occurs before engulfment, at the stage of bacterial attachment.

The attachment defect observed most likely reflects the reduced abundance of membrane-protein transcripts in *Dyrk1a^−/−^* macrophages, although no single receptor emerges, and the large number of affected transcripts makes testing each one impractical. Two candidates stand out because their transcripts are reduced and both have been linked to bacterial uptake: CD36 and SR-B1. CD36 is a broad phagocytic receptor that mediates adhesion and uptake of Gram-positive and Gram-negative bacteria in non-phagocytic cells (52), and its inhibition lowers the proportion of macrophages that internalize *L. monocytogenes* through a Tim3- and Nrf2-dependent mechanism (53). Macrophages lacking SR-B1, encoded by *Scarb1*, internalize fewer *L. monocytogenes* and less dextran, indicating a broad role in phagocytosis (54). Neither receptor has been assessed for a role in adherence, so testing CD36 and SR-B1 in a DYRK1A-dependent manner would show whether either contribute to the phenotypes reported here. Beyond receptor abundance, transcriptional regulation by DYRK1A of actin dynamics and Src-family kinase activity could alter receptor distribution and clustering, reducing responsiveness as described for the Fc receptor (55). Alternatively, loss of DYRK1A reduces cell size (56), which would lower membrane surface availability for bacterial attachment.

The reduced abundance of phagocytosis, endocytosis, and immune-receptor transcripts led us to hypothesize that the role of DYRK1A extends beyond *L. monocytogenes*. *Dyrk1a-*deficient macrophages showed reduced internalization of antibody-opsonized beads through the Fc receptor pathway, a route unrelated to *L. monocytogenes* recognition, positioning DYRK1A as a nonspecific regulator of phagocytosis. Testing a panel of bacterial species, we found that the closely related *L. ivanovii* and *L. innocua*, as well as the more distant *B. subtilis*, strongly require DYRK1A for internalization. DYRK1A also contributed to *S. aureus* internalization, though more modestly than for the other Gram-positive species. *S. aureus* enters macrophages through multiple receptors (57,58), so a DYRK1A-independent route may partially compensate, although this remains to be tested. The requirement was least clear for the Gram-negative *P. aeruginosa*, where intracellular CFU were not significantly changed, as measured by gentamicin protection assay, but significantly fewer *Dyrk1a-*deficient macrophages internalized at least one bacterium as measured by pHrodo. The pHrodo signal can arise from dead bacterial membranes, and *P. aeruginosa* is rapidly killed in the phagosome (59), so we regard the requirement of DYRK1A for Gram-negative uptake as tentative. Overall, DYRK1A is required for internalization of multiple Gram-positive species and for Fc-mediated phagocytosis, with a possible but less certain contribution to Gram-negative uptake.

Pathway analysis of the *Dyrk1a^−/−^* transcriptome also identified gene sets for immune receptor and cytokine response pathways. A role for DYRK1A in immune signaling was expected from prior work. Recent studies have also identified a role for DYRK1A in the TLR4/NFκB signaling axis, which drives expression of IL-8, IL-6, IL-1β, Type II IFN, and TNF-α, all clusters identified in our macrophage transcriptome (38,44). The prior studies linking DYRK1A to TLR4/NFκB signaling were performed in neuroinflammation models. Our observations show that DYRK1A also contributes to immune signaling in macrophages, adding innate immunity to the documented roles of the kinase in the central nervous system and adaptive immunity. The breadth of downregulated innate immunity transcripts is notable, indicating that transcriptional shifts driven by DYRK1A may be a mechanism of broad immune regulation. This may be especially relevant during infection, since LPS stimulation increases DYRK1A protein (44). DYRK1A may also shape immune responses through direct substrate phosphorylation, a non-transcriptional mechanism not examined here.

Our results identify DYRK1A as a broad regulator of bacterial uptake by macrophages, a step of infection studied far less than the intracellular killing that follows. These findings open several questions for future investigation. Whether the phagocytic role of DYRK1A requires catalytic activity remains open, particularly since DYRK1A supports coronavirus entry independently of kinase activity (39). A kinase-dead rescue would distinguish a catalytic from a scaffolding function. Extending these findings to primary macrophages and animal models will test whether the role of DYRK1A in phagocytosis influences the outcome of infection *in vivo*. Growth-permissive macrophages can promote *L. monocytogenes* dissemination, so defining how DYRK1A shapes bacterial uptake may reveal how the balance between spread and clearance is set during listeriosis. More broadly, these findings expand the role of DYRK1A from the nervous system to innate immune defense, establishing macrophage phagocytosis as a new arena for DYRK1A biology.

## METHODS

### Bacterial strains and culture conditions

All bacterial strains used in this study are summarized in Table S3. Bacteria were grown on either brain heart infusion (BHI) agar (*Listeria spp.)* or Luria-Bertani (LB) agar (*Bacillus subtilis*, *Staphylococcus aureus*, *Pseudomonas aeruginosa*, *Escherichia coli*) at 37°C overnight unless otherwise stated. Bacteria were cultured in their respective broth medium overnight while shaking (220 rpm) at 37°C prior to experiments, unless otherwise indicated. Antibiotics (Sigma Aldrich) were used at the following concentrations: streptomycin, 200 µg/mL; chloramphenicol, 7.5 µg/mL; carbenicillin, 100 µg/mL. Plasmids were introduced into chemically competent *E. coli* by heat shock.

### Mammalian cell lines

Cas9-expressing immortalized bone marrow derived macrophages (iBMMs) have been described previously (60). HEK293T cells were purchased from ATCC. Cell lines were incubated at 37°C in 5% CO_2_ in high glucose (4.5 g/L) Dulbecco’s Modified Eagles Medium (DMEM; Gibco) with 10% heat-inactivated fetal bovine serum (FBS; Cytiva) supplemented with sodium pyruvate (2mM) and L-glutamine (1 mM). Cells were maintained in medium containing Pen-Strep (100 U/mL) but plated in antibiotic-free medium 24 hours prior to experiments. Antibiotics were used at the following concentrations: gentamicin, 50 µg/mL; puromycin, 2.5 µg/mL.

### Construction of knockout cell lines

Two individual sgRNAs targeting *Dyrk1b* were designed based on those in the Brie library and were individually cloned into lentiCRISPR_v2 (Addgene #52961) (61,62). Guides were annealed, phosphorylated with T4 Polynucleotide Kinase (Fisher Scientific), then digested and ligated with Esp3I (Fisher Scientific) into lentiCRISPR_v2. The lentiCRISPR_v2 plasmids were transformed into Stbl *E. coli* at 30°C and sequences confirmed via both Sanger (Azenta) and whole plasmid (Plasmidsaurus) sequencing.

Plasmids either containing sgRNA targeting *Dyrk1b* or an empty vector control were transfected into HEK293T cells for lentiviral packaging. Third generation lentiviral packaging plasmids pVSV-G (Addgene #138479), pMDLg/pRRE (Addgene #12251), pRSV-Rev (Addgene #12253) and the lentiviral vector were combined with 1X Polyethylenamine (Fisher Scientific) and added to HEK293T cells. 24 hours post-transfection, cell culture medium was replaced with medium containing 25 mM HEPES. 48 hours post-transfection, supernatant was collected, spun at 300 x *g* for 8 minutes to collect cell debris, and frozen at −80°C. Cas9-expressing iBMMs were transduced by plating cells with transfection supernatants and 8 µg/mL polybrene (Sigma). Beginning 48 hours post-transduction, cell lines were passaged under antibiotic selection with puromycin for 10 days. To validate genetic editing, genomic DNA was isolated from cell lines via Quick-DNA Miniprep Kit (Zymo) and cut sites were amplified and Sanger sequenced. Sequencing was analyzed via Interference of CRISPR Edits (ICE; EditCo) to approximate knockout efficiency (31).

*Dyrk1a-*deficient cell lines were previously generated using an analogous protocol (10). *Dyrk1a* single cell clonal populations were generated via single-cell sorting on BDFACS Aria 2 from both *Dyrk1a* sgRNA1 and sgRNA 2 knockout populations.

### Intracellular growth curves

The day before infection, iBMMs were plated at a density of 5 × 10^5^ per well in 24-well TC-treated dishes (Corning). Cultures of *L. monocytogenes* were inoculated from single colonies grown on agar and cultured at 30°C, stationary overnight. Bacteria were washed twice with sterile phosphate-buffered saline (1X PBS) and resuspended in pre-warmed iBMM medium. iBMMs were infected at a multiplicity of infection (MOI) of 1 and incubated at 37°C in 5% CO_2_. After 30 minutes of infection, cells were washed twice with 1X PBS and medium containing gentamicin was added to kill extracellular bacteria. At 0.5, 2, and 5 hours post-infection (p.i.), cells were washed twice with 1X PBS and lysed in 250 µL cold 0.1% TritonX-100. Lysates were serially diluted and 10 µL drips were plated to enumerate intracellular CFU.

### Gentamicin protection assay

Intracellular bacteria were measured by gentamicin protection assay, as previously described (63,64). In brief, iBMMs were plated at a density of 5 × 10^5^ cells per well into 24-well TC-treated dishes 24 hours prior to infection. Bacterial cultures were inoculated from single colonies grown on agar into either BHI (*Listeria spp.*) or LB (*B. subtilis, S. aureus, P. aeruginosa*) and cultured overnight, shaking at 37°C. Bacteria were then diluted to an OD_600_ appropriate for growth to mid-log over 3 hours while shaking at 37°C (0.01 *B. subtilis*, *S. aureus*; 0.05 *L. monocytogenes*, *P. aeruginosa*; 0.10 *L. ivanovii*; 0.15 *L. innocua*). Bacteria were washed three times with 1X PBS and diluted in pre-warmed cell culture medium to concentrations corresponding to MOI 0.01, 0.1, 1, 10, or 100 per 1 mL. iBMM medium was replaced with 1 mL of bacteria-containing medium, and infected cells were spun at 300 x *g* for 3 minutes to synchronize bacterial contact with cells. Cells were then incubated at 37°C in 5% CO_2_. 30 minutes p.i., cells were washed twice with 1X PBS and medium containing gentamicin was added to kill extracellular bacteria. After 1 hour of infection, cells were washed twice with 1X PBS and lysed in 250 µL cold 0.1% TritonX-100. Lysates were serially diluted and 10 µL drips were plated on LB agar for enumeration of intracellular CFU.

### Internalization assay by flow cytometry

Internalization of pHrodo-labeled bacteria was measured as previously described, with the following alterations (65). Briefly, iBMMs and bacteria were plated and cultured as described in the section above. Prior to infection, bacteria were diluted to an OD_600_ of 0.01-0.15 and grown three hours to mid-log while shaking at 37°C. Bacteria were washed twice with 1X PBS and diluted to an OD_600_ of 1 in 1X PBS. Bacteria were pelleted, resuspended in 200 µL of 1 µM pHrodo Red (Thermo Scientific) in 0.1 M sodium bicarbonate, pH 9.3 and 1% DMSO, and incubated at room temperature with rotation for 45 minutes. Labeling was then quenched with pHrodo wash buffer C. Labeled bacteria were washed twice with pHrodo wash buffer C and twice with 1X PBS. Bacteria were diluted to MOI 0.01 – 100 in pre-warmed cell culture medium and iBMMs were infected as described in the above section. 30 minutes p.i., cells were washed twice with 1X PBS and detached from the cell culture dish with EDTA-Trypsin (0.05%) for 5 minutes at 37°C in 5% CO_2_. Detached cells were treated with FACS Buffer (2 mM EDTA and 5% FBS in 1X PBS) to deactivate trypsin and transferred to FACS tubes on ice. Samples were analyzed on a FACSymphony A1 (BD Biosciences) flow cytometer. Phagocytes were gated based on their forward scatter (FSC-A) and side scatter (SSC-A), and doublets were omitted based on their forward scatter area (FSC-A) and forward scatter height (FSC-H). Internalization was measured by comparing forward scatter (FSC-A) to PE fluorescence (PE-A) to quantify pHrodo-positive events in comparison to total events with the use of FlowJo.

### Immunoblotting

iBMMs were seeded in T75 flasks for 24 hours before cells were lysed in cold RIPA Buffer (150 mM NaCl, 200 mM NP-40, 0.5% sodium deoxycholate, 0.1% SDS, 50 uM Tris, pH 8) containing Halt Protease and Phosphatase Inhibitor Cocktail (Thermo Scientific). Lysates were incubated on ice for 20 minutes, then centrifuged at 20,000 x *g* at 4°C for 20 minutes to pellet cellular debris. Supernatants were collected, flash frozen, and stored at −80°C. Lysates were thawed on ice and protein concentration quantified by Pierce^TM^ BCA Protein Assay Kit (Thermo Scientific). Total protein (40 ng) was separated by SDS-PAGE on a 7% Acrylamide gel and transferred onto PVDF membrane (BioRad) via wet transfer at 100V over 1 hour. Membranes were blocked for 30 minutes with Intercept (TBS) Blocking Buffer (LI-COR). Membranes were then incubated at 4°C rocking overnight in antibody cocktail containing anti-DYRK1A (Abnova H00001859-M01 lot# P7211-7D10) diluted 1:1,000 and anti-myosin IIa (Cell Signaling Technologies #3403S) diluted 1:1,000 in Antibody buffer (50% Intercept (TBS) Blocking Buffer, 50% TBST). The next day, membranes were washed in TBST and incubated for 1 hour rocking at room temperature with 1:5,000 goat anti-rabbit Alexa Flour 680 (Invitrogen) and 1:5,000 goat anti-mouse Alexa Flour 800CW (LI-COR) in Antibody buffer. Membranes were washed three times in TBST, then twice in PBS before imaging on a LiCor Odyssey CLx.

### Immunofluorescence microscopy

iBMMs were plated on sterile 12mm round glass coverslips (Mercedes Scientific) in 24-well plates at a density of 5 × 10^5^ cells per well. *L. monocytogenes* constitutively expressing *mCherry* were incubated overnight in BHI at 37°C while shaking. Prior to infection, bacteria were diluted to an OD_600_ of 0.05 in BHI and incubated at 37°C, with shaking, for three hours to mid-log. Bacteria were washed twice in 1X PBS and diluted in pre-warmed cell culture medium. Cells were infected with an MOI of 2 or 5, centrifuged at 300 x *g* for 3 min to synchronize bacterial contact with cells, then incubated at 37°C in 5% CO_2_. To achieve countable infection dynamics, cells infected with MOI 5 were assessed at 15 minutes p.i., and cells infected with MOI 2 were assessed at 30 or 60 minutes p.i.. At each timepoint, cells were washed twice with 1X PBS and coverslips were removed and inverted onto 25 µL of 4% formaldehyde (Thermo Scientific) on parafilm for 10 minutes for fixation. Coverslips were washed 30 times by dipping into three 50 mL tubes of 1X PBS, 10 times each. This washing method was performed after each subsequent staining. To prevent non-specific antibody binding to phagocyte Fc receptors, coverslips were incubated in staining buffer (1% BSA in 1X PBS) for 10 min at 4°C, then inverted onto 25 µL Fc block (BD Biosciences) diluted in 1:50 in Antibody buffer for 30 minutes. For immunofluorescence staining, coverslips were incubated with 1:100 *Listeria O* antisera (BD Biosciences) in staining buffer for 30 minutes, then stained with 1:200 goat anti-rabbit AlexaFlour488 (Invitrogen) in staining buffer for 30 minutes in the dark. Stained coverslips were mounted by inversion onto glass slides with ProLong Diamond Antifade Mountant with DAPI (Invitrogen) and allowed to dry overnight. Slides were imaged within one week on a Keyence BZ-X710 inverted fluorescence phase contract microscope at 100X magnification, and images were analyzed using BZ-X Analyzer and ImageJ.

### Phagocytosis of IgG-opsonized fluorescent beads

iBMMs were seeded at a density of 5 × 10^5^ cells per well into one well of a 24-well plate per genotype. Phagocytosis of IgG-opsonized fluorescent latex beads was assessed using IgG FITC Phagocytosis Assay Kit (Cayman Chemical) according to manufacturer instructions. In brief, beads were diluted 1:200 in pre-warmed cell culture medium and 500 µL was incubated with iBMMs for 30 minutes. Cells were washed in Phagocytosis Assay Buffer, and quenched with 0.04% trypan blue for 2 minutes, then washed again in Phagocytosis Assay Buffer. Cells were detached from the cell culture dish with EDTA-Trypsin (0.05%) for 5 minutes at 37°C in 5% CO_2_, quenched in Flow buffer (5% FBS in PBS), and transferred to FACS tubes on ice. Samples were analyzed on a LSRII (BD Biosciences) flow cytometer. Phagocytes were identified by forward scatter (FSC-A) and side scatter gating (SSC-A), and doublets were omitted based on their forward scatter area (FCS-A) and forward scatter height (FSC-H). FITC+ cells were measured by comparing FITC fluorescence to a no-bead control at 5% background using FlowJo.

### RNA isolation and sequencing

Cells of each genotype were plated at a density of 5 × 10^5^ cells per well into three wells of a 24-well plate and incubated at 37°C with 5% CO_2_. 24 hours after plating, cells were washed with 1X PBS then lysed in 350 µL total LPB lysis buffer (Macherey-Nagel RNA isolation NucleoSpin® RNA Plus Kit) and flash frozen for storage at −80°C. Lysates were thawed and RNA isolated according to Macherey-Nagel RNA isolation NucleoSpin RNA Plus Kit. Isolated RNA was quantified via nanodrop and stored at −80°C. Total RNA was added to reaction buffer from the SMART-Seq v4 Ultra Low Input RNA Kit for Sequencing (Takara), and reverse transcription was performed followed by PCR amplification to generate full length amplified cDNA. Sequencing libraries were constructed using the NexteraXT DNA sample preparation kit with unique dual indexes (Illumina) to generate Illumina-compatible barcoded libraries. Libraries were pooled and quantified using a Qubit Fluorometer (Life Technologies). Sequencing of pooled libraries was carried out on a NextSeq 2000 sequencer (Illumina) with paired-end 59-base reads, using a NextSeq 2000 P4 XLEAP SBS Reagent Kit (Illumina) with a target depth of 5 million reads per sample. Base calls were processed to FASTQs on BaseSpace (Illumina), and a base call quality-trimming step was applied to remove low-confidence base calls from the ends of reads. The FASTQs were aligned to the GRCm38 mouse reference genome, using STAR v.2.4.2a and gene counts were generated using htseq-count. QC and metrics analysis was performed using the Picard family of tools (v1.134). Pathway enrichment analysis was performed as previously described using GSEA, Cytoscape and EnrichmentMap (35).

## ACKNOWLEDGEMENTS

We would like to acknowledge the Benaroya Research Institute Genomics Core (RRID:SCR_026658) for RNA-sequencing. The M.J. Murdock Charitable Trust generously supported the equipment used by this core facility. We are grateful to Lauren Shull, Lieselotte Kreuk, and Rochelle Glover for valuable discussions. This work was supported in part by the Howard Hughes Medical Institute.

