## Supplemental Materials for "Macrophage DYRK1A is broadly required for bacterial phagocytosis"

#### SUPPLEMENTAL MATERIAL

Figure S1. DYRK1A loss broadly alters the macrophage transcriptome

Figure S2. DYRK1A contributes to macrophage uptake of laboratory and clinical *S. aureus* strains

Table S1. RNA-seq (Separate Excel File)

Table S2. GSEA Pathway Enrichment Analysis output (Separate Excel File)

Table S3. Strains used in this study

Table S4. Oligonucleotides used in this study

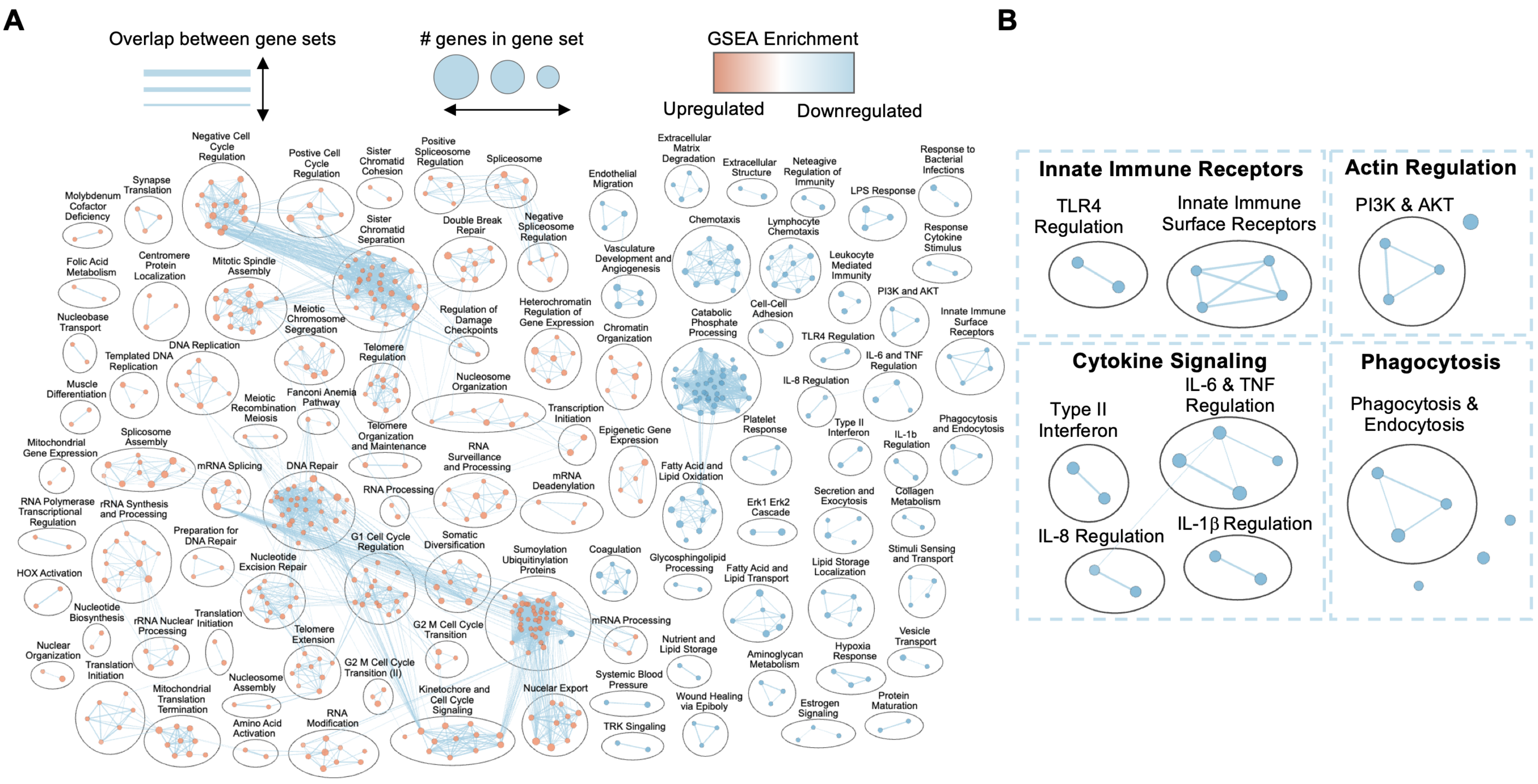
A
B
Overlap between gene sets
### genes in gene set
GSEA Enrichment
Upregulated
Downregulated
Negative Cell Cycle Regulation
Postive Cell Cycle Regulation
Sister Chromatid Cohesion
Positive Spliceosome Regulation
Spliceosome
Synapse Translation
Molybdenum Cofactor Deficiency
Double Break Repair
Negative Spliceosome Regulation
Sister Chromatid Separation
Folic Acid Metabolism
Centromere Protein Localization
Mitotic Spindle Assembly
Meiotic Chromosome Segregation
Regulation of Damage Checkpoints
Heterochromatin Regulation of Gene Expression
Nucleobase Transport
DNA Replication
Templated DNA Replication
Telomere Regulation
Nucleosome Organization
Chromatin Organization
Muscle Differentiation
Meiotic Recombination Meiosis
Fanconi Anemia Pathway
Telomere Organization and Maintenance
Transcription Initiation
Mitochondrial Gene Expression
Splicosome Assembly
mRNA Splicing
DNA Repair
RNA Surveillance and Processing
mRNA Deadenylation
Epigenetic Gene Expression
RNA Polymerase Transcriptional Regulation
rRNA Synthesis and Processing
Preparation for DNA Repair
RNA Processing
G1 Cell Cycle Regulation
Somatic Diversification
Sumoylation Ubiquitinylation Proteins
Coagulation
HOX Activation
Nucleotide Excision Repair
Nucleotide Biosynthesis
rRNA Nuclear Processing
Translation Initiation
Telomere Extension
G2 M Cell Cycle Transition
mRNA Processing
Nuclear Organization
Translation Initiation
Nucleosome Assembly
G2 M Cell Cycle Transition (II)
Kinetochore and Cell Cycle Signaling
Nucelar Export
Mitochondrial Translation Termination
Amino Acid Activation
RNA Modification
Systemic Blood Pressure
TRK Singaling
Extracellular Matrix Degradation
Extracellular Structure
Neteagive Regulation of Immunity
LPS Response
Response to Bacterial Infections
Endothelial Migration
Chemotaxis
Lymphocyte Chemotaxis
Response Cytokine Stimulus
Vasculature Development and Angiogenesis
Leukocyte Mediated Immunity
PI3K and AKT
Cell-Cell Adhesion
Catabolic Phosphate Processing
TLR4 Regulation
IL-6 and TNF Regulation
Innate Immune Surface Receptors
IL-8 Regulation
Type II Interferon
Platelet Response
IL-1b Regulation
Phagocytosis and Endocytosis
Fatty Acid and Lipid Oxidation
Erk1 Erk2 Cascade
Secretion and Exocytosis
Collagen Metabolism
Glycosphingolipid Processing
Fatty Acid and Lipid Transport
Lipid Storage Localization
Stimuli Sensing and Transport
Nutrient and Lipid Storage
Aminoglycan Metabolism
Hypoxia Response
Vesicle Transport
Wound Healing via Epiboly
Estrogen Signaling
Protein Maturation
Innate Immune Receptors
TLR4 Regulation
Innate Immune Surface Receptors
Actin Regulation
PI3K & AKT
Cytokine Signaling
Type II Interferon
IL-6 & TNF Regulation
IL-8 Regulation
IL-1β Regulation
Phagocytosis
Phagocytosis & Endocytosis

**Figure S1. DYRK1A loss broadly alters the macrophage transcriptome.** (A) Enrichment map of gene sets differentially enriched between *Dyrk1a*$^{-/-}$ and EV macrophages by GSEA, visualized in Cytoscape with the EnrichmentMap plugin [1]. Nodes are gene sets (size, number of genes; color, normalized enrichment score, with red upregulated and blue downregulated); edges connect gene sets that share genes. Gene sets shown meet FDR q <0.05 using MSigDB Database. (B) Magnified view of the immune and phagocytosis clusters, with representative genes of decreased transcript abundance grouped by functional category.

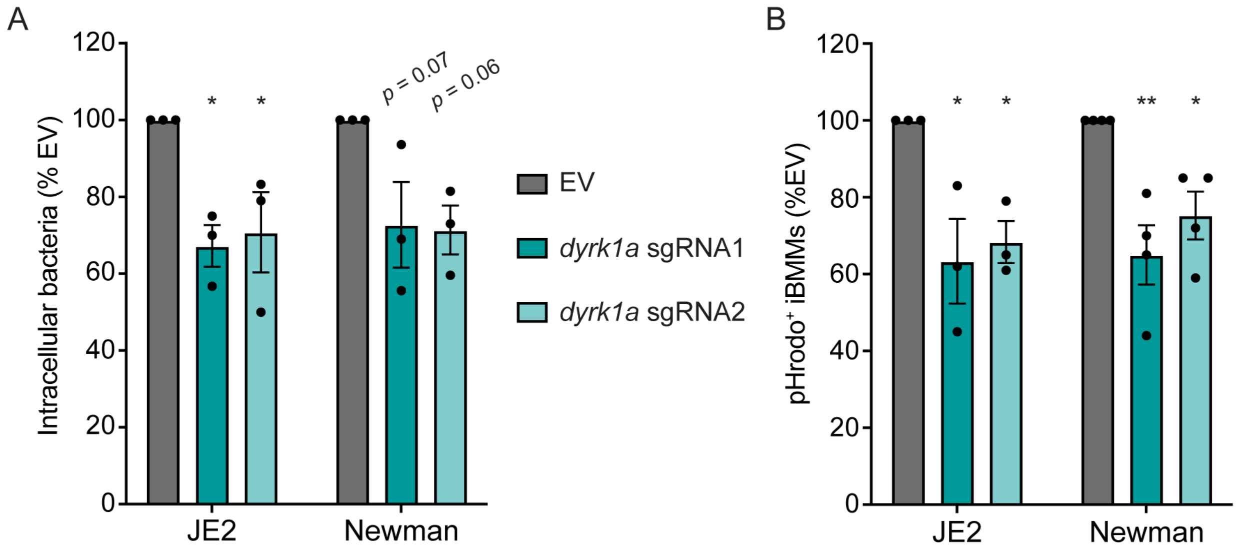


**Figure S2. DYRK1A contributes to macrophage uptake of laboratory and clinical *S. aureus* strains.** (A) Gentamicin protection assay quantifying intracellular *S. aureus* 1-hour p.i. in DYRK1A-deficient and EV iBMMs, for the clinical MRSA strain JE2 and the laboratory strain Newman (MOI = 0.001). Each data point is the mean of two technical replicates. (B) pHrodo phagocytosis assay quantifying the proportion of iBMMs containing pHrodo-labeled *S. aureus* by flow cytometry at 30 minutes p.i., for JE2 and Newman (MOI = 0.1). (A-B) Bars are means and SEM of at least three biological replicates, normalized to EV iBMMs. Exact *p* values are shown for comparisons with $p > 0.05$; *$p < 0.05$, **$p < 0.01$, as determined by Dunnett's multiple comparisons test compared to EV in each condition.

**Table S3. Bacterial strains used in this study**

| Name | Species | Strain | Description | Source or Reference |
|---|---|---|---|---|
| | *E. coli* | Stbl | for strain construction | NEB |
| MPK0982 | *E. coli* | Stbl | lentiCRISPR.*Dyrk1B*-sgRNA1 | This work |
| MPK0983 | *E. coli* | Stbl | lentiCRISPR.*Dyrk1B*-sgRNA2 | This work |
| MLR-E1102 | *E. coli* | XL1 | lentiCRISPR.*Dyrk1A*-sgRNA1 | [2] |
| MLR-E1103 | *E. coli* | XL1 | lentiCRISPR.*Dyrk1A*-sgRNA2 | [2] |
| MLR-L0001 | *L. monocytogenes* | 10403S | | [3] |
| MLR-L0551 | *L. monocytogenes* | 10403S | pPL2.pHyper-mCherry | [4] |
| MLR-L0516 | *L. ivanovii* | | | [5] |
| MLR-L0432 | *L. innocua* | CLIP11262 | | [6] |
| MLR-B0704 | *B. subtilis* | MB4 | hly::Sp-β | [7] |
| MLR-S0004 | *S. aureus* | Newman | | [8] |
| MLR-S0858 | *S. aureus* | JE2 | | [9] |
| AK74 | *P. aeruginosa* | PA01-WZ | | [10] |

**Table S4. Oligonucleotides used in this study**

| Name | Sequence (5'-3') | Description | Reference |
|---|---|---|---|
| dyrk1b_sgRNA1_ICE_F | ATGCCCCAGAGCATGCAGGGTA | Amplify *dyrk1a* sgRNA1 site for ICE analysis | [2] |
| dyrk1b_sgRNA1_ICE_R | AAGCTCAGCTGACAGCAAGGGC | | [2] |
| dyrk1b_sgRNA2_ICE_F | AAATGGTGCGGTGCTGGGGATG | Amplify *dyrk1a* sgRNA2 site for ICE analysis | [2] |
| dyrk1b_sgRNA2_ICE_R | AGGGAGAGGCTGCTTCCCTGAG | | [2] |
| dyrk1b_sgRNA1_F | CACCGGTTGTCGTCATCATAACCG | *Dyrk1b* sgRNA2 from Brie library | [11] |
| dyrk1b_sgRNA1_R | AAACCGGTTATGATGACGACAACC | | [11] |
| dyrk1b_sgRNA2_F | CACCGCACCCCGAAAGTGTGTGTTG | *Dyrk1b* sgRNA3 from Brie library | [11] |
| dyrk1b_sgRNA2_R | AAACCAACACACACTTTCGGGGTGC | | [11] |
| dyrk1b_sgRNA1_ICE_F | CGGTGGACCTCATCAAGACC | Amplify *Dyrk1b* sgRNA1 site for ICE analysis | This study |
| dyrk1b_sgRNA1_ICE_R | ACACAATCTGGCCCTCAGTG | | This study |
| dyrk1b_sgRNA2_ICE_F | GGAAGGTGTGAGAGACGCAA | Amplify *Dyrk1b* sgRNA2 site for ICE analysis | This study |
| dyrk1b_sgRNA2_ICE_R | GGGAGGCTCTGAGCAATGTT | | This study |

**Supporting Information References:**
